# Live cell GLUT4 translocation assay reveals *Per3* as a novel regulator of circadian insulin sensitivity in skeletal muscle cells

**DOI:** 10.1101/2025.02.13.638110

**Authors:** Rashmi Sivasengh, Andrew Scott, Brendan M. Gabriel

## Abstract

Type 2 Diabetes (T2D) is a growing global health concern, with skeletal muscle playing a central role in its pathology due to its role in postprandial glucose disposal. Insulin resistance in skeletal muscle often precedes T2D diagnosis and is linked to impaired GLUT4 trafficking. Additionally, circadian disruptions are increasingly recognized as contributors to metabolic dysfunction. We have shown that skeletal muscle cells from T2D donors exhibit disrupted CLOCK/BMAL1 rhythms, impairing mitochondrial oxidative function. To investigate circadian insulin sensitivity, we developed a high-throughput, live-cell assay for GLUT4 translocation, allowing us to explore novel regulators of metabolic rhythmicity in skeletal muscle.

In this study, we analysed circadian insulin sensitivity in skeletal muscle using transcriptomics, PheWAS, and GLUT4-HiBiT assays. L6 and human myocytes were cultured, transfected, and assessed via our novel live-cell, circadian GLUT4 translocation assay. Circadian rhythms were analysed (JTK_CYCLE), and siRNA knockdowns targeted key genes, with glucose uptake measured using the Uptake-Glo assay in human myocytes.

Skeletal muscle cells from individuals with T2D exhibited disrupted circadian rhythms, with altered rhythmicity in *ARNTL, HOXB5, PER3*, and *TSSK6* (*p*<0.05). Public database analysis revealed significant associations between these genes and T2D (*p*=0.0001–0.03). Our high-sensitivity, live-cell assay demonstrated a 120% increase in GLUT4 translocation at 40 nM insulin (*p*<0.001). Circadian rhythmicity in GLUT4 translocation was evident at physiological insulin levels (5 & 30 nM). *PER3/Per3* knockdown significantly reduced GLUT4 translocation (*p*<0.001) and glucose uptake in human muscle cells (*p*<0.01), while *Arntl* knockdown caused a moderate reduction (*p*<0.01) in GLUT4 translocation. *Per3* silencing also abolished circadian rhythmicity in GLUT4 translocation (*p*<0.01).

These findings establish *PER3/Per3* as a key regulator of circadian GLUT4 translocation and insulin sensitivity, linking its dysregulation to metabolic impairments in T2D. Additionally, this study presents a novel live-cell GLUT4 translocation assay that is highly sensitive and capable of performing time-course or circadian experiments in the same group of cells. The sensitivity of this assay, coupled with reduced variation that is possible from using live-cells in time-course assays make it a powerful new tool for the discovery of GLUT4 translocation regulators. In this study, we have identified *Per3* as a novel regulator of GLUT4 translocation in skeletal muscle cells.

## 1. Introduction

Type 2 Diabetes (T2D) is a growing, global threat to health, placing a high burden on healthcare services. Skeletal muscle is a central organ within the pathology of T2D, being the major storage depot for postprandial glucose disposal (Gabriel and Zierath, 2017). Insulin resistance of skeletal muscle is recognized as a key driver of T2D disease progression (DeFronzo and Tripathy, 2009). In fact, the changes to metabolic pathways which ultimately lead to insulin resistance of this tissue may be present years before other signs of the development of T2D can be detected (Gabriel and Zierath, 2017). For example, induced pluripotent stem cells from donors with T2D that have been differentiated into myoblasts have multiple defects, including reduced insulin-stimulated glucose uptake and reduced mitochondrial oxidation (Batista et al., 2020). These T2D-associated defects are conserved despite the transformations that these cells undergo during experimental procedures. It has not been fully elucidated how cultured myocytes from T2D donors preserve a dysfunctional phenotype, including insulin resistance (Bouzakri and Zierath, 2007) but likely reflects genetic background and epigenetic mechanisms. One key molecular defect that appears to underpin insulin resistance in skeletal muscle is defective trafficking of the glucose transporter GLUT4 to the cell membrane in response to insulin signaling in a temporally appropriate manner. Indeed, insulin resistance impedes the ability of peripheral tissue to respond flexibly to metabolic events throughout the day and people with T2D have disrupted circadian metabolism (Gabriel and Zierath, 2022). It is known that altered sleep/wake rhythms from shiftwork, sleep disorders, and social jet lag are associated with obesity, T2D, and related (Mokhlesi et al., 2019; Shan et al., 2018; Tan et al., 2018; Vetter et al., 2018), emphasising the role of circadian rhythms in metabolic health. Physiological circadian rhythms are ultimately driven by cell autonomous circadian rhythms (Panda, 2016). These are generated by a transcription-translation autoregulatory feedback loop composed of transcriptional activators CLOCK and BMAL1 (ARNTL) and their target genes Period (*PER*), Cryptochrome (*CRY*), and REV-ERBα (*NR1D1*), which rhythmically assemble to complete a repressor complex that interacts with CLOCK and BMAL1 to eventually inhibit transcription (Takahashi, 2015). We have shown that cell autonomous rhythms of *CLOCK* and *BMAL1* are disrupted in skeletal muscle cells from donors with T2D and that this is coupled with a loss of mitochondrial oxidative circadian rhythm (Gabriel et al., 2021). This loss of mitochondrial functional rhythm was linked with a reduction in circadian transcripts associated with the inner-mitochondrial membrane in cells from people with T2D. Further, *in vivo* inner-mitochondrial gene expression was highly associated with whole-body insulin sensitivity when participants underwent a hyperinsulinaemic-euglycaemic clamp and skeletal muscle biopsies (Gabriel et al., 2021). This led us to hypothesise that a loss of circadian rhythmicity in specific metabolic targets was causative in the reduction of skeletal muscle insulin sensitivity. A key factor in skeletal muscle’s ability to respond to insulin and uptake blood glucose is the insulin-stimulated activation of GLUT4 (Gabriel and Zierath, 2017). However, methods to analyse circadian insulin sensitivity in terms of GLUT4 trafficking in skeletal muscle have until now relied on endpoint assays (Heckmann et al., 2022) which reduces throughput and introduces experimental variability into the assay. We have developed a high-sensitivity, high-throughput, live-cell, circadian assay to measure GLUT4 translocation in skeletal muscles cells. With this new tool, we are able to fully test our hypothesis and identify novel regulators of circadian insulin sensitivity in skeletal muscle.

## 2. Materials and Methods

### 2.1 Identifying novel regulators for circadian insulin sensitivity in skeletal muscle

In this study, initial data preprocessing involved the application of the ‘**filterByExpr** function from the ‘**edgeR** package, which was utilised to selectively exclude features with low expression levels, thereby enhancing the robustness of the dataset. Subsequently, the ‘**removeBatchEffect** function from the ‘**limma** package was employed to correct for any potential batch effects, ensuring that observed variations were attributable to biological differences rather than technical artifacts. Post-preprocessing, the data were converted to log Counts Per Million (logCPM) values for each detected feature, facilitating a standardised comparison across samples. This processed dataset, including all features, is publicly accessible via the NCBI repository at https://www.ncbi.nlm.nih.gov under the GEO Accession code GSE182117. For analytical purposes, the data were extracted and normalized using the base R scale function. This normalization process transformed the data into Z-scores, thereby standardizing them for more effective comparative analyses and visualization. Phenome-wide association study (PheWAS) analysis was conducted using the ATLAS PheWAS database (https://atlas.ctglab.nl/PheWAS). Genetic variants associated with diabetes-related traits were queried, and statistical associations with multiple phenotypes were retrieved.

### 2.2 Cell culture conditions and compounds

L6-GLUT4 HiBiT cells were grown in MEMα supplemented with 10% fetal bovine serum (FBS) (PAA, Tet free) and 2.5 µg/mL blasticidin (Invitrogen). L6-GLUT4 HiBiT cells were incubated in starvation media 24 hours prior to each experiment and serum shocked for circadian experiments. For differentiation, the cells were grown in MEMα + supplemented with 2% horse serum (Lonza) for 7 days. HSMM (human skeletal muscle myoblasts) cells were grown in Gibco™ DMEM/F-12, HEPES supplemented with human epidermal growth factor (hEGF) (3 µg/mL), dexamethasone (0.1%) and, FBS (10%). For differentiation, the cells were grown in Gibco™ DMEM/F-12, HEPES + supplemented with 2% horse serum (Lonza) for 7 days. All cells were grown at 37°C and 5% CO2.

### 2.3 Cell line creation – transfection of GLUT4-HiBiT

In this study, we developed a stable myocyte cell line that expresses GLUT4-HiBiT in skeletal muscle cells. Our choice of L6 Rat skeletal muscle cells (ATCC CRL-1458) was guided by their high insulin sensitivity, making them particularly suitable for insulin sensitivity assays (Abdelmoez et al., 2020). The expression vector, TK-HiBiT-GLUT4, was provided by Promega UK limited. We followed the transfection procedure outlined below (Prosen et al., 2013). For the selection of stable clones, we applied 2.5 µg/mL of blasticidin (Invitrogen). **Magnetofection:** for each transfection, 4 µg of DNA was diluted in 200 µL serum free media. Prior to each use, the PolyMag or PolyMag Neo tube was vigorously vortexed. 15 µL of PolyMag was added to a microtube or a U-bottom microwell. The 200 µL of DNA solution was then immediately mixed with the PolyMag or PolyMag Neo solution through vigorous pipetting. The mixture was incubated for 20 minutes, allowing for the formation of the transfection complex. The transfection mix (DNA + PolyMag or PolyMag Neo) was then added to the cells in T25 flask. The cell culture plate (T25) was placed upon a magnetic plate for a duration of 20 minutes to facilitate transfection. **Stable cell line generation (Post Transfection):** cells were incubated at 37°C in a CO2 incubator for 48 hours to allow for the expression of the HiBiT vector. After 48 hours, the medium was replaced with fresh culture medium containing 1.0 µg/mL (Lower Concentration) blasticidin (Invitrogen). The medium containing blasticidin was replaced every 2-3 days. Cells were monitored daily, and the selection process was continued with an increase in the concentration of blasticidin until colonies of blasticidin-resistant cells were visible, typically within 2-3 weeks. Stable clones were screened for the expression of HiBiT using Nano Glow HiBiT Assay.

### 2.4 Insulin dose response in myotubes using Nano-Glow HiBiT assay

L6-GLUT4-HiBiT cells were resuspended to 5^*^1000 cells in 100 µL of growth medium, plated in 96 well Greiner Flat Transparent plates. After myotubes formation, cells were serum starved for 24 hours. On the day of the experiment, the cells were washed twice with PBS, and 100 µl of serum free MEMα media was added with 5 mM glucose (Sigma, filter sterilized). The cells were treated with gradient dose of insulin (0, 5, 10, 20, 30, 40) (stock: 1 mg/mL) and immediately screened. Translocation of GLUT4 from intracellular storage pool to plasma membrane was quantified using Nano-Glow HiBiT detection tool from Promega UK Limited. An equal volume of Nano-Glo HiBiT Reagent (100 µL) (Promega N3030), consisting of Nano-Glo HiBiT Buffer, Nano-Glo HiBiT Substrate, and LgBiT Protein, were added according to the manufacturer’s protocol, and cells were incubated for 15 minutes at room temperature with shaking. Luminescence was measured using a Tecan Spark® 10M plate reader (Tecan, Männedorf) controlled by SparkControl™ V2.0 software with 0.5 s of integration time in room temperature. Data were normalized to the value of untransfected cells (no GLUT4-HiBiT).

### 2.5 Real time luminescence recording in myotubes – 60 hours

After postfusion, myotubes were synchronized with serum shock (50% FBS + MEMα, supplemented with 2.5 µg/mL Blasticidin) for 2 hours in 37°C in a cell culture incubator, then the medium was changed to 100 µl of serum-free MEM α media with 5 mM glucose (Sigma, filter sterilized) to create the low-glucose condition. Luminescence was measured every 6 hours for over 60 hours period using Nano-Glo -HiBiT detection assay. Fresh media was replenished at the end of every reading. Luminescence was normalized to the value of untransfected cells (No GLUT4-HiBiT). Error bar, SD (*n*=3).

### 2.6 Rhythmicity analysis

Rhythmicity was assessed using JTK cycle using R. JTK_CYCLE. The analysis was conducted using the ‘JTK_CYCLEv3.1’ R script. The R (v4.2.1) environment was set to handle strings as non-factors. Two data files, “x.annot.txt’ and ‘x.txt’, were imported. The search for circadian rhythms was narrowed to periods of 60 hours, corresponding to 5 time points per cycle. This was set using the **periods** variable. The results were converted to a data frame and then subjected to Benjamini-Hochberg (BH) correction for multiple testing using the **p.adjust** function. The corrected P-values (BH. Q), alongside the JTK_CYCLE output (ADJ.P, PER, LAG, AMP), were collated with the annotation data. The results were sorted based on adjusted P-value and amplitude. The scripts used are publicly available: https://github.com/RashmiSivasengh/circadian-rhythm.

### 2.7 siRNA mediated knockdown and real time polymerized chain reaction

Commercially available siRNA targeting PER3 (Human cells) / Per3, Arntl, Tssk6 and Hoxb5 were purchased from Thermo Fisher Scientific. L6 skeletal muscle myocytes were cultured with MEM α and 10% fetal bovine serum (FBS). Cells were maintained at 37 °C in a humidified atmosphere with 5% CO2. Cells were plated at 60 to 80% confluence using trypsin-EDTA (Gibco, Thermo Fisher Scientific). Cells were differentiated at 90% confluency and transfected at day 01 of differentiation. L6 cells were transfected using Lipofectamine RNAiMAX (Invitrogen, Thermo Fisher Scientific) following the manufacturer’s protocol (Invitrogen, 2020). A stock solution of siRNA was prepared at a concentration of 40 µM. For each well of a 6-well plate, 60 µL of OPTI-MEM (Gibco, Thermo Fisher Scientific) was mixed with 3.6 µL of Lipofectamine RNAiMAX. In a separate tube, 50 µL of OPTI-MEM was combined with 5 µL of siRNA, resulting in a final siRNA concentration of 200 pmoles/L. After gentle mixing, 55 µL from the first tube containing the Lipofectamine RNAiMAX mixture was added to the siRNA solution. The combined mixture was then incubated at room temperature for 15 minutes to allow the formation of siRNA-lipid complexes. Meanwhile, the cells were washed with phosphate-buffered saline (PBS), and 1 mL of AMEM supplemented with 2% horse serum was added to each well. Subsequently, 100 µL of the siRNA final mixture was directly added to the cells, which were then incubated for 12 hours. Following incubation, the media was replaced with fresh differentiation media. The cells were allowed to stabilize for 24 hours before conducting further assays. qPCR was carried out for determining gene expression by using Fast SYBR Green Master Mix (Thermo Fisher Scientific) and predesigned TaqMan Gene Expression Assays (Thermo Fisher Scientific). TaqMan rat housekeeping genes: B2m.

### 2.8 Glucose uptake of Human skeletal muscle myotubes

The assay was performed using the glucose Uptake-Glo TM Assay kit (Promega corporation), which provides a luminescent readout based on the detection of 2–deoxy glucose–6 phosphates (2DG6P), a glucose analog, as an indicator of glucose uptake activity. Human cells were plated in a 96 well plate at a density of 10,000 cells per well and allowed to adhere overnight. Post fusion, *PER3* gene was silenced, cells were serum-starved for 24 hours and followed by glucose uptake assays.

## Results

### 3.1 Disrupted circadian rhythms in skeletal muscle cells from individuals with T2D

Re-analysis of publicly available data (Gabriel et al., 2021) demonstrated that *ARNTL, HOXB5, PER3*, and *TSSK6* had a loss of rhythmicity or a differential circadian rhythmicity in skeletal muscle cells from people with T2D compared to healthy individuals matched for age and BMI [Figure 1]. Our analysis of publicly available data revealed *ARNTL, HOXB5, PER3*, and *TSSK6* were all significantly associated with T2D across a variety of databases [Table 1]

**Table 1:** Significant genes identified from public database.

| Gene | atlas ID | PMID | Year | Domain | Trait | P-Value | N |
| --- | --- | --- | --- | --- | --- | --- | --- |
| *ARNTL* | 4045 | 30054458 | 2018 | Endocrine | Type 2 Diabetes | 0.003 | 659256 |
| *HOXB5* | 4083 | 28566273 | 2017 | Endocrine | Type 2 Diabetes | 0.0001 | 159208 |
| *PER3* | 4176 | 30718926 | 2019 | Endocrine | Type 2 Diabetes | 0.014 | 191764 |
| *TSSK6* | 4176 | 30718926 | 2019 | Endocrine | Type 2 Diabetes | 0.03 | 191764 |

**Figure 1.**
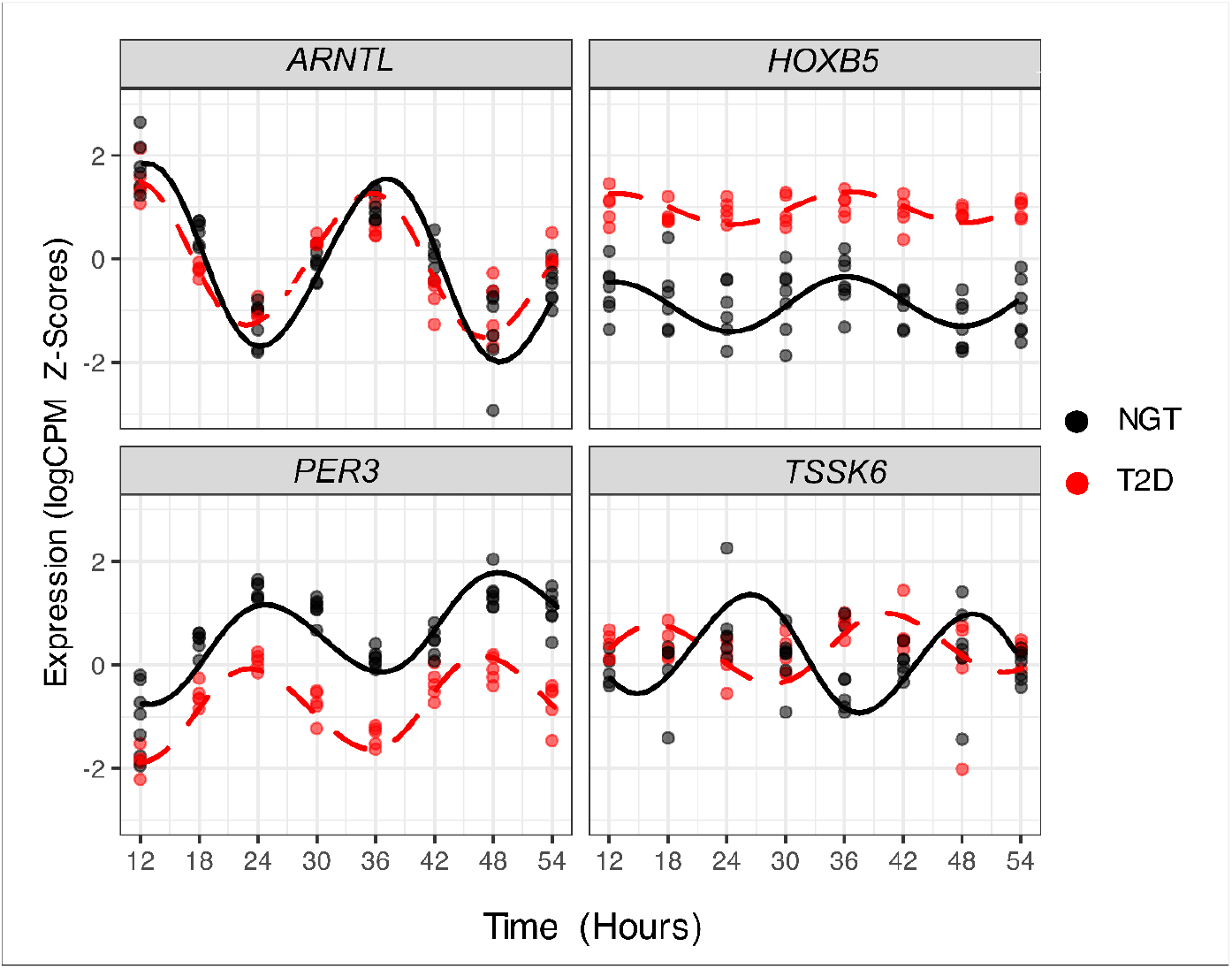
Disrupted circadian gene expression in myocytes from people with T2D: **A**. Circadian rhythmicity in gene expression, comparing T2D (shown in red) with Normal Glucose Tolerance (NGT, in black). The z-score of logCPM is plotted against time since synchronization, illustrating gene expression patterns over a 24-hour cycle. Harmonic regressions fitted to the data highlight the circadian trends in both groups. The comparison emphasises the distinct circadian gene expression profiles between T2D and NGT individuals.

### 3.2 Live-cell GLUT4 translocation assay: model validation

To evaluate the utility of our novel system for monitoring circadian insulin sensitivity, we developed stable skeletal muscle cell lines expressing HiBiT-tagged GLUT4 (Figure 2A-representing the plasmid map). Quantification of GLUT4-HiBiT expression in these clones was performed using the Nano-Glo HiBiT Assay as depicted in Figure 2B. The assay detected significant luminescence in a subset of clones transfected with 4000 ng of DNA, with mean luminescence values exceeding the baseline established by non-transfected controls (NT). The observed luminescence levels were approximately 1000-fold (Figure 2C) higher than controls, confirming specific expression of the HiBiT-tagged GLUT4 protein. We next conducted an insulin dose-response experiment using L6 cells transfected with the GLUT4-HiBiT construct. The resulting data exhibited a linear increase of luminescence with increasing insulin concentration. Cells expressing the GLUT4-HiBiT construct exhibited a substantial increase in luminescence, approximately 120-fold (Figure 2D) higher compared to uninduced cells (0 nM insulin). The luminescent response reached its maximum at an insulin concentration of 40 nM, indicating saturation of the GLUT4 translocation mechanism at this dose. These findings demonstrate that L6 cell-derived myotubes possess a high degree of insulin sensitivity, evidenced by their robust luminescent response. Moreover, the assay displayed excellent throughput and sensitivity, providing an effective means to quantify GLUT4 translocation in skeletal muscle-derived cells.

**Figure 2:**
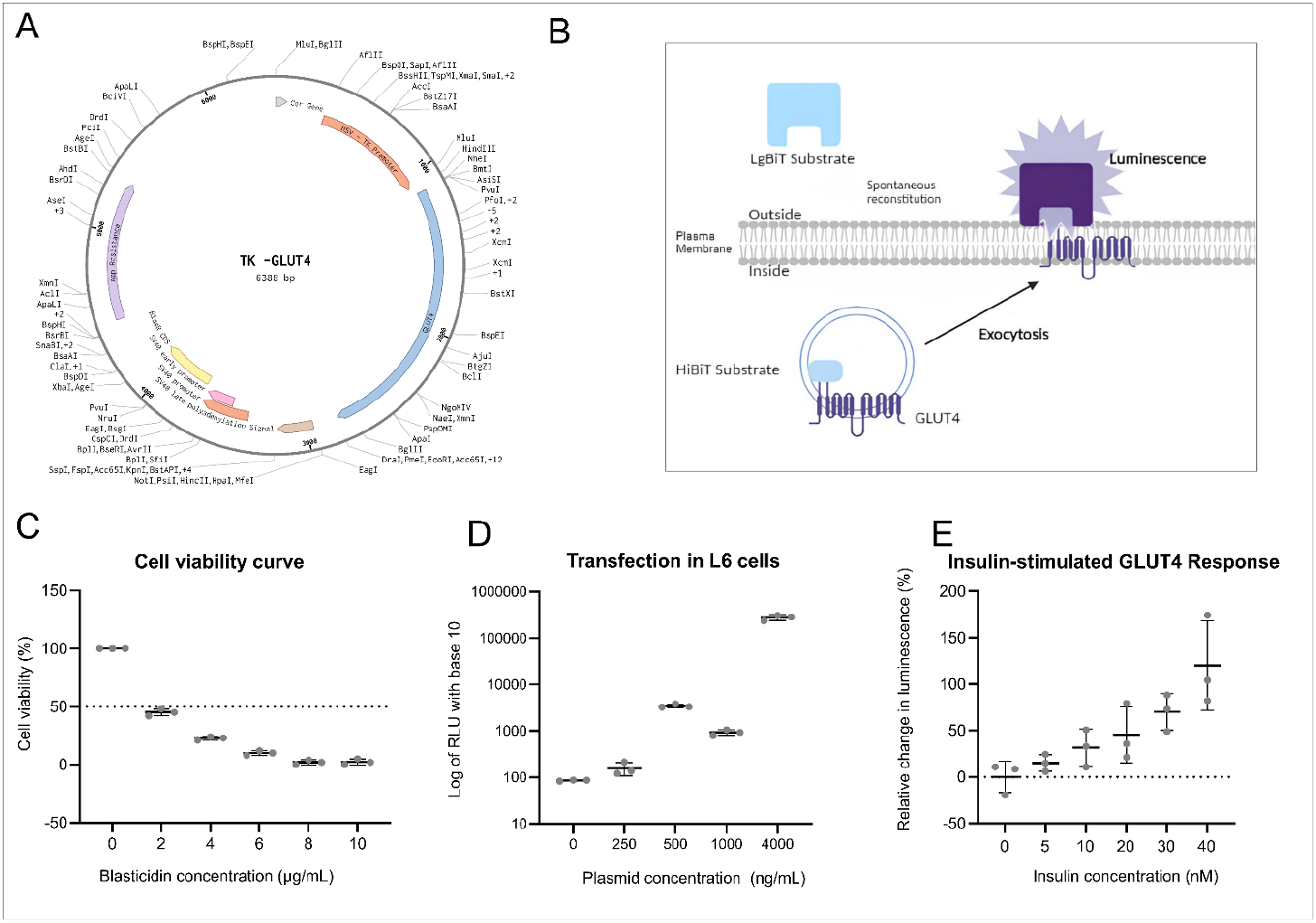
Validation of live-cell GLUT4 translocation assay. Luminescence detection of GLUT4-HibiT exocytosis upon insulin stimulation. (A) Systematic representation of plasmid map of HiBiT expression vector. (B) Schematic representation of luminescence detection of surface GLUT4 upon exocytosis using spontaneous reconstitution of HiBiT and LgBiT substrate. (C) Kill curve of skeletal muscle cells (L6 cells) with varying concentration of antibiotic blasticidin. (D) Luminescence detection of entire GLUT4 in cytoplasm after lysis of the cells, across various concentrations of DNA used for transfection. Luminescence showing 10^4 fold increase normalized to non-transfected cells (NT). Error bars, *n*=3. (E) Luminescence measurement upon various concentrations of insulin. Error bars, *n*=3.

### 3.3 Testing circadian dose-dependent GLUT4 translocation response to insulin

To evaluate the impact of different insulin concentration (0 nM to 50 nM) on circadian rhythmicity in gene expression, JTK analysis was conducted on time-course data. The analysis revealed rhythmicity was significant only at 5 nM and 30 nM insulin concentrations (Figure 3A), with both conditions exhibiting a consistent 24-hour period. Other concentrations lacked statistically significant rhythms, suggesting a dose-dependent disruption or dampening of circadian control (Table 2). Due to the homogeneity of response and the greater amplitude of rhythmicity, 30 nM insulin, which is close to a physiological insulin concentration, was selected for further experiments.

**Table 2:** Overview of the JTK analysis for different insulin concentrations.

| Concentration (nM) | BH. Q | Adj.P | PER | LAG | AMP |
| --- | --- | --- | --- | --- | --- |
| 5 | 0.002 | **0.002** | 24 | 17 | 1.42e<sup>-14</sup> |
| 10 | 1 | 1 | 24 | 12 | 2.49e<sup>-14</sup> |
| 20 | 1 | 1 | 24 | 6 | 1.78e<sup>-14</sup> |
| **30** | 0.003 | **0.003** | 24 | 9.5 | 184.57 |
| 40 | 0.53 | 0.53 | 24 | 8.5 | 1.07e<sup>-14</sup> |
| 50 | 0.3 | 0.3 | 24 | NA | NA |

**Figure 3:**
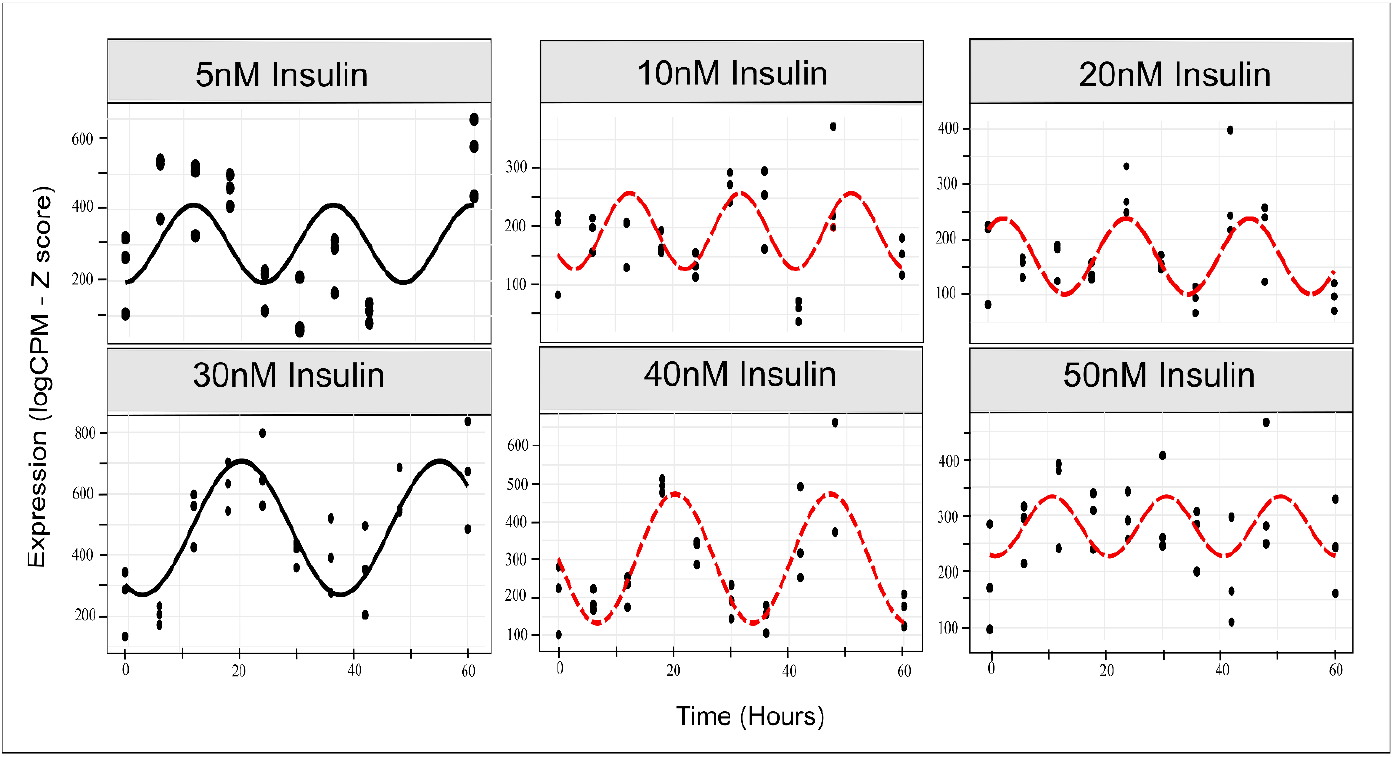
Circadian rhythm of GLUT4 translocation. **A** The circadian patterns of GLUT4 translocation in L6 skeletal muscle cells across varying insulin concentrations, analysed over a 60-hour time. The data plotted using R (v4.2.1) with a harmonic regression script.

### 3.4 Per3 Knockdown Impairs GLUT4 Translocation in L6 Cells Following siRNA-Mediated Gene Silencing

To investigate the role of regulatory genes in GLUT4 translocation, siRNA-mediated knockdown was performed targeting *Per3, Tssk6, Arntl, and Hoxb5* in L6 cells. The efficiency of gene knockdown was validated using RT-qPCR (Figure 4A). Significant reductions in relative mRNA expression were observed for *Per3, Tssk6*, and *Arntl (p*<0.01) in response to silencing. Next, the effect of gene silencing on GLUT4 translocation was assessed under insulin stimulation (30 nM). As shown in Figure 4B, knockdown of *Per3* resulted in a significant reduction in GLUT4 translocation, measured as luminescence relative to control cells (*p*<0.001). Similar reductions were observed following *Arntl* knockdown (*p*< 0.01), whereas *Tssk6* knockdown showed no significant effect (*p>*0.05). These results highlight *Per3* and *Arntl* as novel regulators of GLUT4 trafficking in skeletal muscle cells.

**Figure 4:**
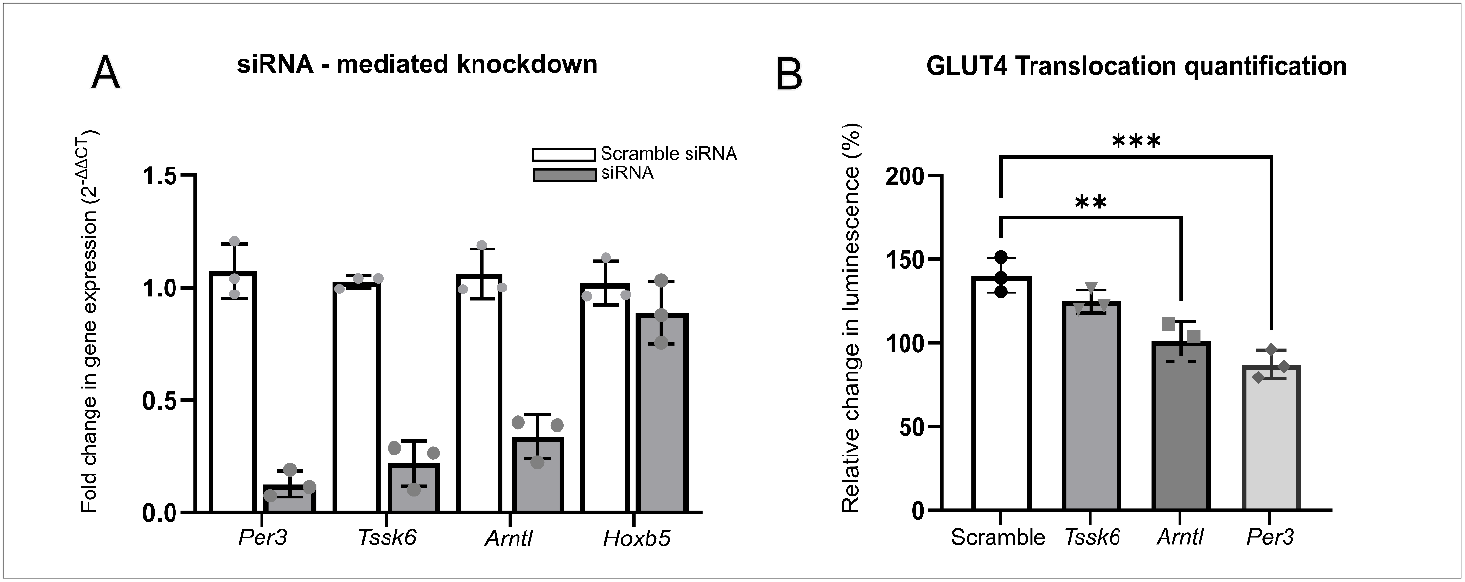
Effect of target gene knockdown on GLUT4 Translocation. (A) Validation of siRNA-mediated knockdown efficiency for *Per3, Tssk6, Arntl*, and *Hoxb5* genes in L6 cells. Relative gene expression was quantified using qPCR and normalized to control (untreated cells). Data are presented as mean ± SEM (Standard Error of the Mean) (*n*=3 biological replicates). Significant knockdown was observed for *Per3, Tssk6*, and *Arntl* compared to control (*p*<0.01). (B) Quantification of GLUT4 translocation in response to 30 nM insulin stimulation following siRNA knockdown of *Per3, Tssk6*, and *Arntl*. Translocation was measured as the relative change in luminescence compared to control cells. Knockdown of *Per3* and *Arntl* significantly reduced GLUT4 translocation (^***^*p*<0.001, ^**^*p*<0.01), while *Tssk6* knockdown caused a moderate but statistically non-significant reduction (*p*>0.05). Data are presented as mean ± SEM (*n*=3 biological replicates).

### 3.5 Per3 knockdown disrupted GLUT4 translocation in a circadian rhythm-dependent manner in skeletal muscle cells

Given that *Per3* displayed the most robust reduction of GLUT4 translocation, we selected this as a target to study in a circadian time-course assay. Time-course analysis revealed that GLUT4 translocation was disrupted in *Per3* knockdown cells compared to scrambled control cells under 30 nM insulin stimulation (Figure 5A). The rhythmic pattern observed in the control group was abolished in *Per3-*deficient cells, suggesting that *Per3* is essential for maintaining the circadian rhythm-dependent regulation of GLUT4 trafficking (Figure 5A). To explore the metabolic consequences of *Per3* knockdown, glucose uptake assays were conducted. In human cells, the knockdown of *PER3* resulted in significantly reduced glucose uptake under insulin-stimulated conditions, compared to non-targeting controls (Figure 5B, *p*<0.01). Collectively, these results highlight the critical role of *Per3*/*PER3* in coordinating circadian rhythm-regulated GLUT4 translocation and metabolic functions, with significant implications for understanding glucose uptake mechanisms in both murine and human systems (Figure 5C).

**Figure 5.**
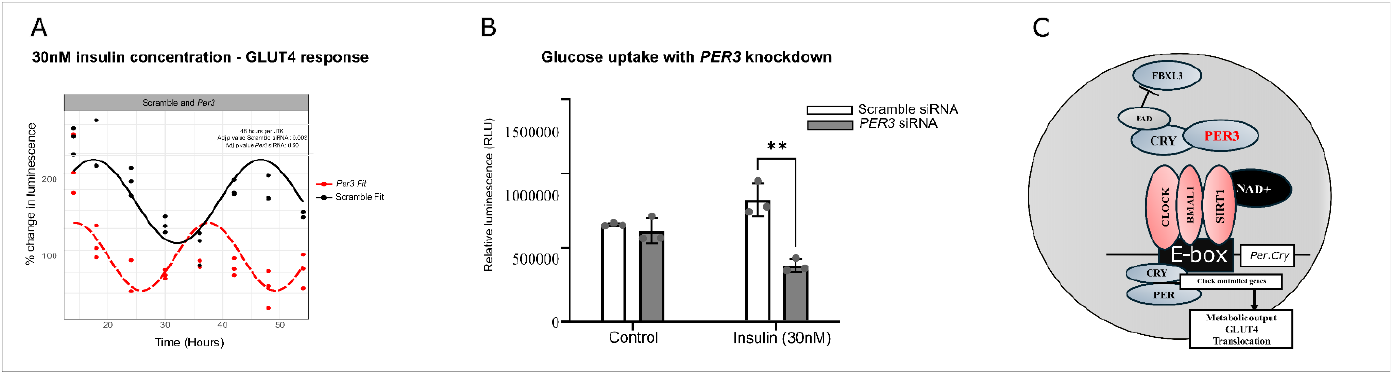
Role of *Per3* in Regulating GLUT4 Translocation and Glucose Uptake. (A) Time-course analysis of GLUT4 translocation under 30 nM insulin stimulation in *Per3* knockdown and scrambled control cells. Data represent the percentage change in luminescence over time, with fitted curves illustrating the distinct patterns of GLUT4 response between *Per3* knockdown (red) and control (black). Statistical significance was determined using a JTK analysis (^**^*p*<0.01) with *Per3* knockdown losing the rhythm. (B) Glucose uptake in human cells with *PER3* knockdown compared to non-targeting (NT) control cells. Cells were treated with or without 30 nM insulin, and glucose uptake was measured as relative luminescence units (RLU). Knockdown of *PER3* significantly reduced glucose uptake under insulin-stimulated conditions (^**^*p*<0.01). Data are presented as mean ± SEM (*n*=3 biological replicates). Statistical analysis was performed using two-way ANOVA. (C) Schematic representation of the molecular pathway involving *PER3*. The diagram highlights the interaction between *PER3*, circadian regulators, and downstream metabolic outputs, including GLUT4 translocation.

## Discussion

We have identified four promising targets that have intrinsically different cycling patterns between skeletal muscle cells from people with T2D and those who are matched for BMI and age but have normal glucose tolerance. These targets were also all associated with T2D in PheWAS studies. To further test the relevance of these targets in the pathology of T2D, we have developed a high-sensitivity, high-throughput, live-cell, circadian assay to measure GLUT4 translocation of skeletal muscle cells. With this new tool we can perform high-throughput circadian assays of novel regulators of circadian insulin sensitivity in skeletal muscle. Our assay revealed *Per3* as a regulator of GLUT4 translocation in response to insulin, regulating glucose uptake, and circadian insulin sensitivity in skeletal muscle cells.

Our study illuminates the relationship between disrupted circadian rhythms and T2D, particularly in the context of skeletal muscle dysfunction. Skeletal muscle’s pivotal role in glucose homeostasis and insulin resistance in T2D is well-established, underscored by persistent defects in insulin-stimulated glucose uptake and mitochondrial oxidation in myocytes obtained from individuals with T2D (Gabriel et al., 2021). A significant aspect of our findings lies in the characterisation of intrinsic differences in cycling patterns between cells derived from individuals with T2D and age- and BMI-matched controls exhibiting normal glucose tolerance. It has previously been demonstrated that there is a relationship between circadian clocks and insulin resistance, a key component of T2D (Stenvers et al., 2018). It has also been shown that induced pluripotent stem cells (iPSCs) derived from T2D donors, when differentiated into myoblasts, exhibit several abnormalities (Batista et al., 2020). These include diminished glucose uptake in response to insulin (Batista et al., 2020). While research has established a link between insulin sensitivity and circadian genes in T2D, there remains a gap in our understanding of the specific genes involved and their precise roles in insulin sensitivity. This is partly due to the previous absence of adequate tools for investigating the circadian GLUT4 response to insulin.

Central to our study is the development of a novel, high-sensitivity, live-cell circadian assay enabling the measurement of GLUT4 translocation in skeletal muscle cells. All current methods for quantifying GLUT4 translocation, including both classical and microscopic approaches, necessitate cell fixation, rendering them as endpoint assays (Heckmann et al., 2022). Notably, among these endpoint assays, the highest observed increase in GLUT4 translocation, up to two-fold, was achieved through a colorimetric method applied to fixed L6 cells (Wang et al., 1998; Wang et al., 2020). In the current study, insulin dose-response experiments revealed a linear increase in GLUT4 translocation with increasing insulin concentrations, with saturation at 40 nM and a maximal translocation response of 120%. Due to the sensitivity of this assay to relatively low insulin concentrations, when compared to similar end-point assays (Wang et al., 1998; Wang et al., 2020), it can use physiological concentrations of insulin as an input signal. For example, both 5 nm and 10 nm insulin concentrations elicited robust responses of 15% and 32% increased GLUT4 translocation (luminescence), respectively.

In this current study, our novel tool also allows for time-course assessment of insulin-responsive GLUT4 translocation in live-cells. Our dose-response circadian experiments revealed that 30 nm insulin was a promising concentration to develop a circadian insulin-sensitivity assay. In order to test the relevance of this approach, we aimed to screen potential regulators of circadian insulin sensitivity in skeletal muscle. We identified these by re-analysing publicly available data (Gabriel et al., 2021), which revealed a loss of rhythmicity or altered circadian rhythmicity in the genes *ARNTL, HOXB5, PER3*, and *TSSK6* in skeletal muscle cells from individuals with T2D compared to healthy controls. These genes were significantly associated with T2D in PheWAS datasets, highlighting their potential role in circadian dysregulation linked to metabolic disease. Gene silencing experiments further established *Per3*, and *Arntl* as regulators of GLUT4 translocation. Interestingly, knockdown of *Per3* resulted in the most pronounced reduction in GLUT4 translocation and was therefore selected as the target to take forward into a circadian GLUT4 translocation assay. However, it should be noted that knockdown efficiency of *Arntl* was lower than *Per3*, which may influence the magnitude of effect on GLUT4 translocation, and *Arntl*/Bmal1 has previously been implicated as a key modulator of glucose tolerance and insulin sensitivity (Schiaffino et al., 2016). Previous research has suggested no obvious circadian disruption in *PER3* rhythmic gene expression in myocytes from participants with T2D and healthy participants (Hansen et al., 2016), albeit with lower statistical power (*n*=3) than the data analysed in the current study (*n*=5-7) (Gabriel et al., 2021). However, in the current study *Per3* ablation impaired circadian rhythmicity, emphasising its essential role in maintaining circadian-regulated insulin sensitivity. This circadian disruption was at a much greater magnitude (Figure 5A) when compared to the circadian gene expression disruption of *Per3* between T2D and NGT (Figure 1). Additionally, glucose uptake assays confirmed that *PER3* knockdown impaired glucose utilisation under insulin-stimulated conditions in human skeletal muscle cells.

While our study suggests a potential link between disrupted circadian rhythms and insulin resistance in skeletal muscle, further investigations are warranted. It is not clear how these data link to glycolytic and mitochondrial metabolism, and disrupted circadian metabolism plays a role in a loss of appropriate temporal response in these pathways (Gabriel et al., 2021). Understanding the precise mechanisms underlying these intrinsic differences in cycling patterns and their implications for insulin sensitivity, particularly within the context of T2D, remains a critical avenue for future research. For example, the mechanism of GLUT4 translocation inhibition is unclear. It is plausible that mitochondrial crosstalk with the core-clock plays a role (Gabriel and Zierath, 2022). Mitochondria play a critical role in regulating Ca^2+^ signaling in skeletal muscle (Gabriel and Zierath, 2022; Ivarsson et al., 2019), and Ca^2+^ promotes GLUT4 exocytosis and reduces its endocytosis (Li et al., 2014). Given the direct regulation of mitochondrial metabolism by the core-clock in skeletal muscle (Lassiter et al., 2018, Gabriel et al., 2021) it is plausible that this regulatory loop is disrupted by *Per3* ablation. Conversely, family member *Per2* is regulated by skeletal muscle contraction in a calcium-dependent manner (Small et al., 2020), suggesting a possible bidirectional feedback loop. Overall, translating these insights into viable therapeutic strategies necessitates a comprehensive grasp of the regulatory networks governing circadian fluctuations in skeletal muscle insulin sensitivity.

In summary, we have identified disrupted circadian rhythms in key genes (*PER3, ARNTL, TSSK6, HOXB5*) in skeletal muscle cells from individuals with T2D. Using our novel live-cell GLUT4-HiBiT assay, we show that *Per3*/*PER3* is essential for maintaining circadian-regulated GLUT4 translocation and insulin sensitivity. Its knockdown abolished rhythmic GLUT4 translocation and reduced glucose uptake in skeletal muscle cells. These findings highlight the link between circadian dysregulation and T2D, particularly in skeletal muscle, and may inform future therapeutic strategies targeting circadian regulation in insulin resistance.

## Acknowledgment

We would like to thank Promega UK for their kind donation of the expression vector (TK-HiBiT-GLUT4), which we have used to create a stable cell line in L6 cells.

## Data Availability

All scripts used for analysis are publicly available: https://github.com/RashmiSivasengh/circadian-rhythm. Raw Data is available: 10.6084/m9.figshare.28409729. GLUT4 HiBiT biological resources are available upon request.

